# Canine Mammary Tumours (CMTs) exploit Mitochondrial Cholesterol for aggressive reprogramming

**DOI:** 10.1101/2024.06.23.599030

**Authors:** Liana Hardy, Brindha Kannan, Manuel Rigon, Genevieve Benton-Hawthorn, Renato L. Previdelli, Iris M. Reichler, Franco Guscetti, Mariusz P. Kowalewski, Michelangelo Campanella

## Abstract

In human breast cancer the mitochondrial translocator protein (TSPO) aids pro-survival cellular response by facilitating the formation of mitochondrial contact sites with the nucleus termed Nucleus Associated Mitochondria (NAM). Here, we show that TSPO positively associates with the aggressiveness of tissues and cells isolated from Canine Mammary Tumours (CMTs). TSPO is also readily upregulated in reprogrammed mammary tumour cells following long-term deprivation of oestrogen or exposure to the endocrine chemotherapeutic (ET) agent Tamoxifen. The latter triggers mitochondrial handling of cholesterol which is facilitated by TSPO whose upregulation reduces susceptibility to Tamoxifen. TSPO binding ligands boost, on the other hand, the efficacy of Tamoxifen and Chemotherapy agents. In aggressive canine mammary tumour cells, TSPO repression impairs the NF-kB pattern thus confirming the pro-survival role of the NAM uncovered in the human counterpart.

Mitochondrial cholesterol handling via TSPO emerges therefore as a signature in the aggressive reprogramming of CMTs thus advancing our understanding of the molecular mechanisms underpinning this pathology. A novel target mechanism to improve bio-marking and therapeutic protocols is here proposed.

## Introduction

Breast cancer is the most frequent malignancy in women which is nonetheless curable in 80% of patients if diagnosed and treated in its early stages[1]. On the contrary, advanced breast carcinomas with distant organ metastases are often incurable due to the limitations of the available therapies. Breast cancer is a highly heterogeneous disease whose molecular features include activation of oestrogen and progesterone receptors, human epidermal growth factor receptor 2 and mutations in breast cancer type 1/2 susceptibility proteins (BRCA1/2), which predispose genetic risk to breast cancer development. Breast cancer treatments differ according to molecular subtype spanning surgery, radiation, and systemic therapies. Among those, hormone, or endocrine-targeted therapy (ET) holds a degree of selectivity which makes them the first line of treatment[1].

In human patients, ET is therefore adopted as neoadjuvant (before surgery) or co-adjuvant therapy (after surgery) and the vastly adopted drug tamoxifen reduces mortality by 31% in oestrogen receptor (ER)-positive breast cancer patients[2]. However, this approach holds limitations as over 50% of advanced ER-positive breast cancers are intrinsically resistant to tamoxifen and about 40% become resistant during treatment[3 4]. Limitations are even greater in Canine Mammary Tumours (CMTs). In this very cohort of patients even though ET is contemplated as treatment, it fails to deliver sufficient benefit whilst in cases leading to a diametral opposite effect which results in a high level of toxicity.

The capacity to manage canine patients affected by mammary tumours is therefore limited and the preventive procedure of spaying[5], does not result in a disease-free animal[6]. Suitable neoadjuvant and co-adjuvant therapeutic options are therefore needed, and a greater understanding of the CMT’s pathophysiology is prodromic to this.

CMTs have been proposed to be a suitable model in comparative studies particularly so in the advanced stages of the disease[7 8]. Several studies have indicated similarities with the human counterpart for *in situ* pathology, and molecular and visual characteristics[7–10]. However, the intrinsic cellular heterogeneity which characterizes CMTs and the ill-defined role of the hormones in the evolution and progression of the conditions cast doubts on the model suitability to inform human cancer physiopathology. The positive correlation between CMTs and E2, established in humans, has not been recapitulated in dogs[11] indicating that more is to be understood on its pathogenesis. The unveiling of novel traits will lead to novel targets to better manage this condition for which mastectomy remains the only effective approach.

In human and mice, mitochondria hold a pathogenic role in the evolution of aggressive breast cancer by resisting the stimuli aimed at altering their permeability to release factors that initiate programmed cell death. Furthermore, by establishing a retro-communication with the nucleus mitochondria propel the expression of adapting genes which preserve proliferative and survival potential[12]. Equally pathogenic is the mitochondrial biosynthesis of pregnenolone, the precursor of steroid hormones, in the development and progression of mammary tumors[13–15]. However, the oncogenic function of this hormone in humans seems not entirely recapitulated in canine patients. Accordingly, the selective ER modulator tamoxifen, which is part of the standard therapy for women with ER-positive breast cancer[16 17], is not recommended in dogs due to its partial agonistic potential and the associated side effects[18 19].

Hitherto nothing is known about the role of mitochondria in CMTs despite this would advance our understanding of intracellular mechanisms which play a part in the genesis and development of the disease.

Over the years we have investigated mitochondrial targets linked with cell pathology and amenable to pharmacological regulation focusing our activity on the 18-kDa translocator protein (TSPO), an outer mitochondrial membrane-based protein ubiquitously expressed in tissues and conserved among species[20 21].

TSPO is involved in the mitochondrial translocation of cholesterol as a part of a larger protein complex, referred to as “transduceosome“[22]. This is a critical step for the generation of steroid precursors representing therefore a regulatory step in the hormonal rewiring at the basis of mammary glands’ malignancy. TSPO works in complex with the steroidogenic acute regulatory protein (STAR) which allows for binding and shuttling of cholesterol into the mitochondrion. TSPO expression positively correlates with an increase in motility, tissue invasion and higher proliferation of malignant mammary neoplasia[23]. Such a correlation is retained in other types of tumours[24].

In human-derived cells of aggressive mammary cancer, we have shown that TSPO-enriched mitochondria form contacts with the nucleus to drive the expression of oncogenes: a communication facilitated by a cholesterol redistribution from the mitochondria in the perinuclear region[25].

TSPO is indeed required for this inter-organellar interaction whereby together with a pool of proteins (ACBD3, PKA, and AKAP95), it can tether mitochondria and nucleus[25]. The pathological remodelling of the mitochondrial network on the nucleus and the consequent evasion of cell death is counteracted by both the downregulation of TSPO as well as its pharmacological inhibition.

In this work, we further corroborate the role of TSPO in CMTs, by appraising its function in the handling of cholesterol and definition of retrograde signalling. Via multiple approaches, we validate the exploitation of the mitochondrial cholesterol pathway as a factor in the aggressive reprogramming of CMTs and its targeting to envisage novel therapies.

## Materials and Methods

### Cell lines

Isogenic cell lines isolated from normal canine mammary glands (CF35), canine mammary adenoma (CF41), and canine mammary carcinoma (REM-134) have been enrolled in the study. CF41 and REM-134 were obtained from ATCC. CF35 cells were donated by Prof. Marilène Paquet University of Montreal, Canada. All canine mammary cell lines were maintained in a 37°C incubator with 5% CO2 in Dulbecco’s modified Eagle Medium (DMEM), supplemented with 10% foetal bovine serum and 100mg/mL penicillin and 100mg/mL streptomycin. To achieve a hormone-deprived environment, CF41 cells were cultured in phenol-red free DMEM, supplemented with 10% charcoal-stripped foetal bovine serum and 1% penicillin/streptomycin. Tamoxifen-resistant CF41 cells (CF41-TAM) were maintained in DMEM supplemented with 10% foetal bovine serum, 100mg/mL penicillin,100mg/mL streptomycin and 5 μM 4-hydroxytamoxifen (H7904 Sigma, UK). For all experiments, cells were seeded to achieve 70% confluency before treatments and transfections.

### Antibodies

Primary antibodies - Anti-PBR/TSPO (Abcam, ab108489); Anti-ERα (ThermoFisher, TA807239); Recombinant Anti-ACBD3 antibody (Abcam, ab134952); Anti-STAR antibody mouse monoclonal (Abcam, ab58013); Anti-NF-κB (ThermoFisher, PA5-16545); Anti-ATPB (Abcam, ab14730). Secondary antibodies - Goat α-rabbit Alexa 488 (Life Technologies, A11008); Goat α-mouse Alexa 594 (Life Technologies, A11032).

### siRNA Transfection

TSPO was transiently knocked down in canine mammary cancer cells using targeted siRNA. TSPO siRNA (Canine)/ Cn2: Target Sequence – TCCTGGTCGCTGAACCTTCCA. Sense: CUGGUCGCUGAACCUUCCATT; anti-sense UGGAAGGUUCAGCGACCAGGA. Cells were seeded in 6-well plate 24 hours before transfection. Cells were transfected with 10 nM canine TSPO siRNA or scrambled siRNA as a control. Transfections were performed using Lipofectamine RNAiMAX as per the manufacturer’s instructions. SiRNA knockdowns were validated for silenced expression using RT-qPCR (**SFig. 2F**).

### Cell viability

Cell viability for dose response was performed using the crystal violet protocol by Feoktistova and colleagues[26]. Propidium iodide staining was used for assessing cell death when crystal violet was not sensitive enough. Quick proliferation assay kits (Abcam 65475) were also used for assessing proliferation.

### Assays of cellular aggressiveness

3D Sphere Formation. Dishes were coated in 1.2% (w/v) poly(2-HEMA) in ethanol and dried in an oven for >5 days at 60°C. Before seeding, plates were placed under UV light for at least 15 minutes to sterilize. Cells were passed through 25G needle to ensure single-cell suspension. 5000 single cells were seeded into each well of a 6-well plate in DMEM/F-12, no phenol-red, no HEPES, with L-glutamine (21041, Gibco) supplemented with B27 and EGF. Spheres were allowed to form for 5 days, before taking measurements of sphere number and diameter. A mammosphere is considered to be ≥50 μm in diameter[27].

Scratch-Wound-Healing assay. Cells were seeded in a 24-well plate and left to grow in 2 ml of DMEM complete media until reaching a full monolayer confluency. On the day of the experiment, the media was removed and replaced with 200 µl of PBS and a p200 pipette tip was used to induce a cross-shaped scratch in each well. After rinsing twice with PBS, 2 ml of fresh DMEM media or DMEM with 200 nM PK11195 was added to each well and four images per well were taken at the four arms of the cross, using the EVOS M7000 imaging system (Thermo Fisher Scientific). The images were taken again at the same positions after 24 hours and then elaborated with the ImageJ software. For the migrating distance quantification, the area not covered by the cells has been measured and subsequentially divided by the length of the scratch on the picture, providing the average distance between the margins. By subtracting the distance between the margins on Day 0 from the distance on DAY 1, we calculated the µm covered by the cells in 24 hours.

### Canine patient samples

RNA and tissue sections from canine patients diagnosed with tumours derived from a previous research[11] were collected at the Vetsuisse Faculty of the University of Zurich, Switzerland. Samples were analysed using qPCR and immunohistochemistry. TSPO, ATP synthase β (ATPB) and β-actin (ACTB) levels of expression were determined using RNA samples of paired healthy and neoplastic mammary tissues from 22 canine subjects. The RNA samples, serum hormone levels, tissue expression of hormone receptors, and reproductive states of the dogs bearing the tumours, were collected, and recorded, and tumours were graded^11^. Paraffin-embedded tissue sections were obtained from Finn’s pathology, canine mammary tissue was previously graded and sorted into the following categories: hyperplasia (non-cancerous/healthy), adenoma (benign), adenocarcinoma (malignant).

### qRT-PCR

RNA extraction was performed on cell pellets using the RNeasy Plus Mini Kit (Qiagen, 74104) following the manufacturer’s instructions. Extracted RNA was quantified and its purity was measured (260/280 nm ratio) using a Spectrophotometer (Nanodrop, Lab Tech. East Sussex, UK). 1 μg of total RNA was used to synthesize cDNA using a QuantiNova Reverse Transcription kit (205410, Qiagen, UK). qPCR was performed using QuantiNova SYBR green (208052, Qiagen, UK) and canine-specific primers (**Table 1**). Samples were run in triplicate using the conditions described in **Table 2**, with biological replicates run on the same plate to avoid batch effects. Expression data was presented either as mRNA copy number when comparing different cell lines or fold-change for isogenic cell lines. The delta-delta Ct (2–ΔΔCt) method was used to quantify and calculate the relative fold gene expression of samples. Gene expression changes in canine patient RNA were presented as fold-change, normalised to gene expression in corresponding healthy RNA from the same patient.

### Cellular fractionation

For the cellular fractionation, we adopted a modified version of a protocol previously described by Operkun and colleagues[28]. Cells were lysed in isotonic buffer A (40 mM Hepes pH7.4, 120 ml KCl, 2 mM EGTA, 0.4% Glycerol, 10 mM β-glycerophosphate and 0.4% NP-40 with protease inhibitors) with the final, critical concentration of the detergent in the lysate equal to 0.2% (approximately 0.3 ml of buffer A for cells growing in 15 cm dish) while rotating for 30 min at 4°C. Nuclei were pelleted by centrifugation at 1000 × g for 5 min and the supernatant was centrifuged further at 10,000 × g for 10 min to obtain the clarified cytosolic fraction. The pellet of nuclei was sequentially and gently washed with 0.1% NP-40 and no-detergent-containing Buffer A and centrifuged at 1000 × g for 5 min; the supernatants were discarded. The nuclear pellet was re-suspended in Buffer B (10 mM Tris-HCl pH 7.4, 1.5 mM KCl, 0.5% Triton X-100; 0.5% Deoxycholate, 2.5 mM MgCl2, with fresh 0.2 M LiCl and protease inhibitors) at a ratio 1:2 v/v and rotated for 1 hour at 4°C and the extract was separated by centrifugation at 2000 × g for 5 min and the supernatant discarded. Pelleted at 2000 × g, the core nuclei were resuspended in 8 M urea, and sonicated to obtain the core Nuclear Fraction.

### Immunoblotting

For the Westen blot analysis, a method similar to what has been previously described has been adopted[29]. Sample proteins were quantified using a BCA Protein Assay Kit (Thermo Fisher Scientific, 13276818). Equal amounts of protein (30 μg) were resolved on 12% SDS– polyacrylamide gel electrophoresis and transferred to 0.45 um pores PVDF membranes (Thermo Fisher Scientific, 88518). The membranes were blocked in 5% nonfat dry milk in TBST [50 mM tris, 150 mM NaCl, 0.05% Tween 20, (pH 7.5)] for 1 hour and then incubated with the appropriate diluted primary antibody at 4°C overnight: TSPO (1:5000; Abcam, ab108489), actin (1:2000; ab8266), acNF-κB (1:500; ab19870) and β-Tubulin (1:1000, Abcam, ab6046). Membranes were washed in TBST (3 × 15 min at RT) and then incubated with corresponding peroxidase-conjugated secondary antibodies (Dako, P0447, P0448) for 1 hour at RT. After further washing in TBST, blots were developed using an ECL Plus on Western Blotting Detection Kit (Thermo Fisher Scientific, 12316992). Immunoreactive bands were analyzed by performing densitometry with ImageJ software.

### Immunohistochemistry

Formalin-fixed paraffin-embedded canine mammary tissue samples of differing tumour classifications (hyperplasia, adenoma, adenocarcinoma) were supplied by Finn’s pathology (n=X/group). Four μm sections were cut using a rotary microtome. Before staining with the appropriate antibodies, samples were de-waxed and rehydrated using xylene and ethanol by standard protocols. Samples were permeabilized using 0.3% triton-x-100 followed by boiling in sodium citrate buffer (10mM sodium citrate, 0.05% Tween 20, pH 6.0) for 10 minutes for heat-mediated antigen retrieval. Tissue was blocked using appropriate goat blocking solution (Abcam, Ab64259) and incubated with primary antibodies: TSPO and ERα diluted in blocking overnight at 4 °C in a humidified chamber. Samples were then washed and incubated with 0.3% H_2_O_2_ for 4 minutes. Following this, samples were incubated for 90 minutes with appropriate secondary pre-diluted antibody from the Vectastain ABC kit (Vector Laboratories Ltd, UK).

Samples were washed in 1X TBS for 10 minutes to remove excess antibodies. Antibody signal was amplified as per instructions using Vectastain ABC reagent. After this, sections were incubated in peroxidase substrate solution (Abcam, Ab64259) until colour was seen. Sections were counterstained with haematoxylin and dehydrated using IMS and Xylene. The sections were then mounted using DPX mounting media. Images were taken using the Leica DM4000 upright bright field microscope on the 40X lens and protein expression was quantified using Image J. The images underwent colour deconvolution using ‘regions of interest’ (ROI) to separate the nuclear (hematoxylin, purple) staining from the protein staining (DAB, red). These separated images were made binary and inverted. A threshold was set and ‘measured’ and the protein staining was normalized to haematoxylin.

### Immunocytochemistry

Cells were grown to 70% confluency on coverslips before fixation in 4% PFA (10 mins, room temp.) followed by three 5-minute washes in PBS. Permeabilization was performed with 0.5% Triton-X in PBS (15 mins, room temp) followed by washing with PBS. Blocking was carried out for 1 hr at room temp in 10% goat serum and 3% BSA in PBS. Primary antibodies were diluted in blocking solution and incubated with cells overnight at 4°C. Cells were washed again before secondary antibody incubation for 1 hr diluted in blocking solution, before a final wash step. Samples were then mounted on slides with DAPI mounting medium (Abcam, ab104139). The following primary antibodies were used: 1:400 ATPB (Abcam ab14730); 1:400 ERα (Abcam, ab32063); 1:200 TSPO (Abcam, ab109497); 1:500, NF-kB (Abcam, ab16502); and the following secondary antibodies: 1:1000 α-mouse Alexa 594 (Life Technologies, A11032); α-1:1000 rabbit Alexa 488 (Life Technologies, A11008).

### Cholesterol measurements

Two methods have been adopted to assess the level of intraorganellar cholesterol. The first one occurred dynamically using Ergosta (dehydroergosterol-ergosta-5,7,9(11),22-tetraen-3β-ol) which was prepared and loaded in the cells as previously described[29]. 20 μg of ergosta was added to the cells in the form of DHE-MβCD complexes and allowed to incubate for 90 minutes at room temperature in 1X HBSS. Before imaging, cells were washed thrice with 1X HBSS and incubated with 20 nM MitoTracker™ Red CMXRos (M7513, Invitrogen). Cells were then washed and imaged in a live imaging solution (A14291DJ, Gibco).

The second one was based on Filipin using a specific kit (Abcam, ab133116) adopted following the manufacturer’s instructions. In this case, too Mitotracker CMXRos (Invitrogen, M7512) was used to stain mitochondria. Images have been visualized with a confocal microscope.

### Confocal Imaging/ImageJ

Fluorescence staining was observed via the Leica SP-5 (Leica Microsystems, Germany) and Zeiss LSM800 (Carl Zeiss AG, Germany) confocal microscopes using oil immersion objectives at 40X and 63X. Images were recorded using Leica-LAS and ZEN softwares. All staining was checked for non-specific antibody labelling using control samples without primary antibody. Confocal imaging was sequential for different fluorophore channels to obtain a series of axial images. All image analysis was done using Fiji (ImageJ; NCBI, USA) and corresponding plug-ins. Quantification of cell protein levels and nuclear to cytosolic ratio was measured by integrated intensity.

### Statistical analysis

Data is presented as mean ± standard error (SEM). Key p-values are indicated on graphs, with a p-value of less than 0.05 determined as statistically significant. Ordinary one-way analysis of variance (ANOVA) using Bonferroni’s test was used for multi-group comparisons. Students t-tests were performed on data to compare means between two datasets. All analyses were performed in GraphPad Prism versions 7 and 8.

## Results

### The cholesterol-binding protein TSPO positively correlates with malignancy of the canine mammary gland

TSPO is part of a complex of proteins which take part in the trafficking of cholesterol from the outer into the inner membrane of mitochondria for the synthesis of pregnenolone[20–22](**Fig. 1 A**). Immunohistochemical analysis of sections from mammary gland hyperplasia (i), adenoma (ii) and adenocarcinoma (iii) isolated from dog patients unveiled a positive correlation between TSPO protein expression level and stage of the disease. An equal correlation was recorded with the protein StAR (**SFig. 1 A, B**) and the oestrogen receptor alpha (ERα) (**Fig. 1 B-D**).

**Figure 1.**
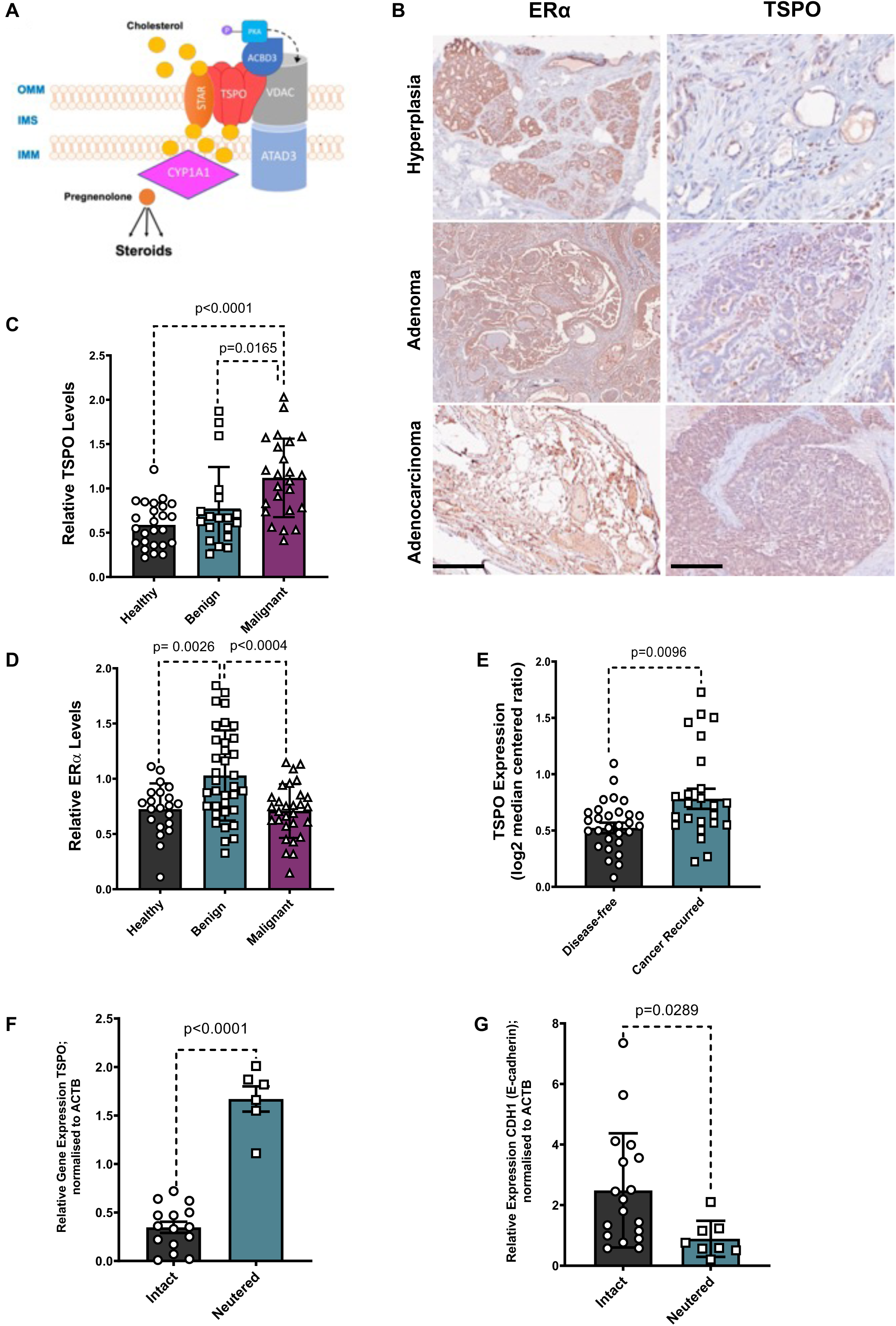
TSPO level of expression positively associates with canine breast cancer aggressiveness. **A)** Schematic representation of the proteins (TSPO, VDAC and STAR) translocating cholesterol from the cytoplasm into the inner mitochondrial membrane. **B)** Immunohistochemical images showing TSPO and ERα expression in canine mammary tissue of different classifications, in order of aggressiveness: Hyperplasia, adenoma and adenocarcinoma. Scale Bar =500 μm. **C)** Quantification of TSPO antigen levels (brown staining) in primary canine mammary tissue shows the highest expression in malignant canine tumours (Healthy 0.8387±0.3373; Benign 1.241±0.3058; Malignant 1.5621±0.6759; n=20). **D)** Quantification of ERα antigen levels (brown staining) which are significantly higher in benign mammary tumours when compared to healthy tissue and malignant mammary cancer in the domestic dog (Healthy 0.9574±0.4928; Benign 1.4379±0.6221; Malignant 0.9517±0.4653; n=20). **E)** Microarray data of TSPO from the Ma Breast 2 study (Ma et al., 2004) shows TSPO expression to be significantly greater in human tumours of which the cancer recurred following tamoxifen treatment. Line at median, error bars show range (Disease-free 0.7797±0.2659; Cancer recurred 1.229±0.3344; n=55, p<0.01). **F)** Fold change expression of TSPO mRNA levels in tissues from neutered and intact dogs with mammary neoplasms. TSPO gene expression is normalised to ACTB (Intact 0.57±0.12; Neutered 1.99±1.35; n=20). **G)** Fold change expression in mRNA of epithelial marker CDH1 (E-cadherin) in neutered and unneutered dogs. CDH1 gene expression is normalised to ACTB (Intact 3.37±1.00; Neutered 1.08±0.35; n=20).

The increased expression of TSPO in the oestrogen (ER) independent stages of the mammary tumours was confirmed in human patients via the analysis of microarray data (**Fig. 1 E**).

Furthermore, the level of TSPO, inspected via quantification of mRNA, was significantly higher in neutered female dogs too (**Fig. 1 F**) in which the expression of the epithelial-mesenchymal transition (EMT) gene CDH1, one of the markers of cancer aggressiveness, was instead reduced[30] (**Fig. 1 G**). To inform the cancer cell biology of TSPO we continued the investigation by enrolling cell lines isolated from benign and malignant tumours of the canine mammary gland.

### TSPO promotes stemness in oestrogen-independent breast cancer cells

We, therefore, assessed the TSPO expression level via immunocytochemistry in the isogenic cell lines isolated from the normal canine mammary gland (CF35), canine mammary adenoma (CF41), and canine mammary carcinoma (REM-134) (**Fig. 2 A-D**) confirming the pattern of expression recorded in tissues. Furthermore, expression analysis of hormone receptors obtained using RT-qPCR highlighted the loss of ESR1 in REM-134 cells (**Fig. 2 E**) in which the % of nuclear ERα was also greater than the corresponding less aggressive counterpart (**SFig. 1 C-E**). Subsequently, the sphere formation potential was assessed in both REM-134 and CF41 cells (**Fig. 2 F, G**). This parameter of the self-renewal capability of cancer cells was greater in the REM-134 cells in which TSPO was prominently overexpressed and this key parameter at the basis of cellular aggressive behaviour was corroborated further via scratch assay analysis (**SFig. 2 A, B**). The TSPO contribution to such a critical feature for malignancy was corroborated further by probing the formation of spheres in the presence of the TSPO ligand PK11195 (acknowledged to act as an antagonist of TSPO in specific conditions) and the compound PMI (p62/SQSTM1-mediated mitophagy inducer)[31 32] which represses TSPO expression whilst promoting mitochondrial quality control via autophagy.

**Figure 2.**
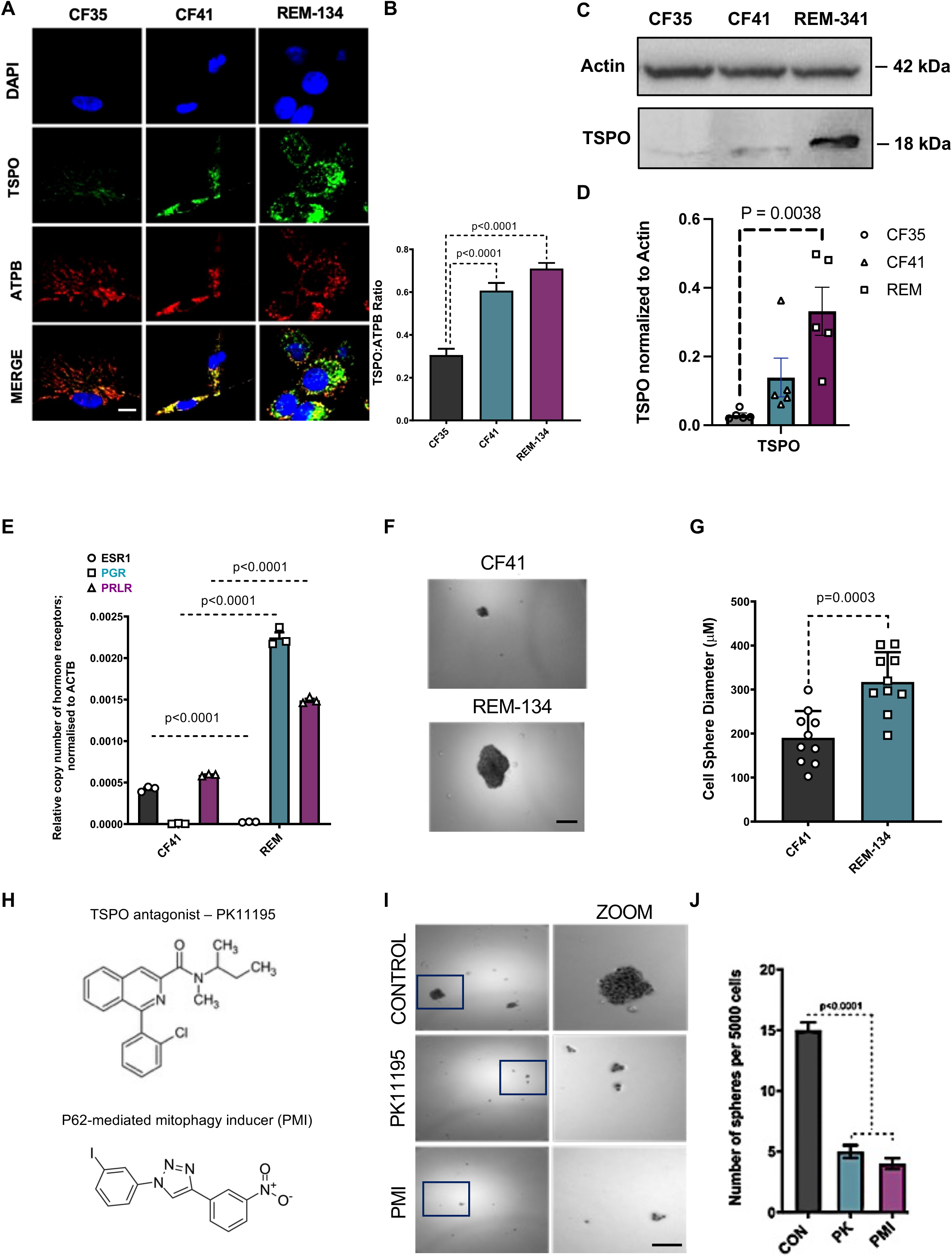
TSPO favours spheroidic growth of Canine Mammary Tumour cells (CMTs). **A)** Confocal images of fluorescent staining of the nucleus (blue), mitochondria (red) and TSPO (green) in canine mammary cell lines. Scale bar = 10 μm. **B**) TSPO protein levels normalised to ATPB quantified using immunofluorescence microscopy (n=15). **C)** Immunoblotting analysis of TSPO in CF35, CF41 and REM-134 with relative quantification reported in **D** (n=3). **E**) Gene expression data showing steroid hormone receptor mRNA copy number in CF41 and REM-134 cells. (CF41 ESR1 4.54E-04±3.92E-04; CF41 PGR 9.09E-06±6.28E-07; CF41 PRLR 6.05E-04±5.79E-04; REM-134 ESR1 2.92E-05±2.12E-05; REM-134 PGR 2.35E-03±2.14E-03; REM-134 PRLR 1.53E-03±1.45E-03; n=3). **F**) Representative images of sphere formation assay using canine mammary cancer cell lines CF41 and REM-134. Scale bar = 200 μm. **G**) Quantification of cell sphere diameter in CF41 and REM-134 cells. 10 spheres > 50 μm in diameter were quantified (CF41 209±171; REM-134 338±295, n=1). **H**) Chemical structures of TSPO ligand PK11195 and mitophagy inducer (PMI). **I**) Representative images of 3D sphere formation assay using CF41 cells treated with DMSO (control), TSPO ligand PK11195, Pravastatin (PVS), mitophagy inducer (PMI) and positive control doxorubicin (DOX). Scale bar = 200 μm. **J)** Quantification of number of spheres per 5000 cells seeded (control DMSO: 15.64±14.36; PK11195: 5.52±4.48; PMI:3.41±2.59; PVS 4.45±3.55; DOX 3.44±2.56; n=6 fields of analysis per condition, p<0.0001).

Both the PK11195[33] (200nM) and PMI (100μM) successfully prevented the formation of spheres in REM-134 cells (**Fig. 2 H-J**).

The different patterns of mitochondrial cholesterol accumulation in response to 4-OH TAM have been further assessed and corroborated via Filipin analysis (**SFig. 2 C, D**).

The ET agent promotes the accumulation of cholesterol in the mitochondria in both cell types analysed even though REM-134 cells feature a higher basal level which we linked to the greater level of TSPO compared to CF41.

The cellular content of NF-kB is also dependent on TSPO whose repression (TSPO KD) alters the pattern and cytosolic availability of this pro-survival transcription factor in REM-134 (**Fig. 3 E-H**) thus recapitulating what was previously recorded in human cells derived from aggressive breast cancer[23].

**Figure 3.**
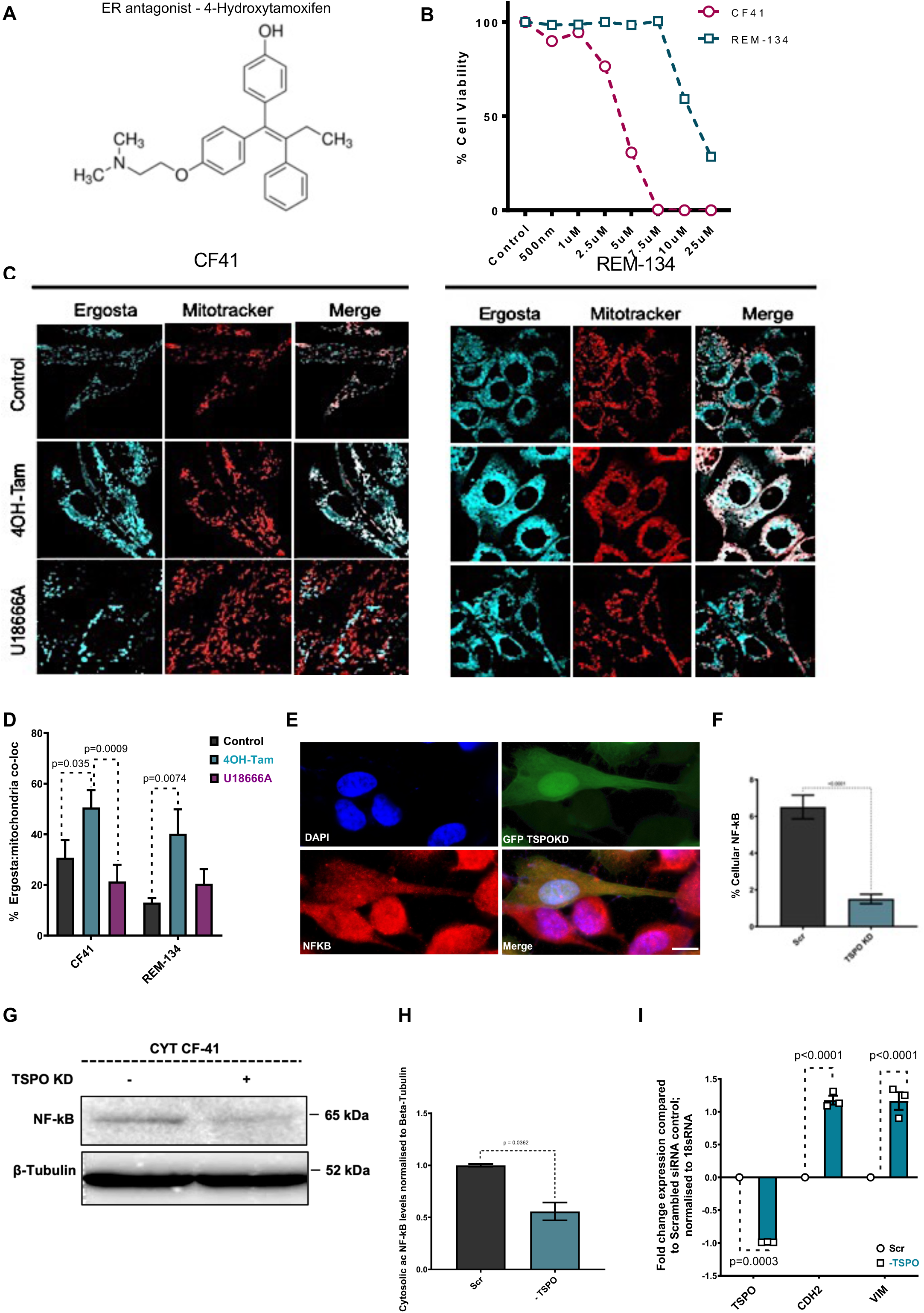
Tamoxifen mobilizes mitochondrial cholesterol in CMTs. **A)** Chemical structure of 4-hydroxytamoxifen. **B**) Quantification of percent cell viability from crystal violet assay according to 4-hydroxytamoxifen concentrations per 14h. **C**) Confocal images of live cells probed for mitochondria (mitotracker; red) and cultured with Ergosta (cyan) following treatments with 4OH-tamoxifen and cholesterol trafficking inhibitor U18666A in CF41 (left) and REM-134 (right) cells. Scale bar = 10 μm. **D**) Quantification of percent Ergosta (cholesterol analogue) co-localising with mitotracker following tamoxifen treatment in CF41 and REM-134 cells. (CF41: Ctrl 37.80±23.74, 4OH-TAM 57.49±43.87, U18666A 27.96±14.88; REM-134 cells Ctrl 22.29±13.08, 4OH-TAM 24.92±21.07; U18666A 18.97±10.90, 15 images were analysed per condition). **E**) Confocal images of CF41 cells co-transfected with GFP (green) and either control siRNA or TSPO knockdown siRNA respectively. Cells were immunoassayed with NF-kB (red). 15 images were analysed per condition. Scale bar = 10 μm. **F**) Quantification of NF-kB level (Scr 24.0±21.5; TSPO KD 15.6±13.1; 15 images were analysed per condition). **G**) Cytosolic enriched fraction of CF-41 control or KD for TSPO immunoblotted for NF-kB and β-Tubulin as control; quantified in **H** (n=3). **I**) Fold change expression of EMT genes –normalised to housekeeping gene 18sRNA- in wild type, REM134 cells and REM134 cells downregulated for TSPO (REMTSPOkd) (TSPO: REM WT=0, REMTSPOkd= -0.99±0.001; CDH2 (N-cadherin): TSPO: REM WT=0, REMTSPOkd= 1.18±0.12; Vim (Vimentin): REM WT=0, REMTSPOkd= 1.19±0.78; n=3).

In addition, TSPO repression (**SFig. 2 E**) impacts the expression of the epithelial-mesenchymal transition (EMT) genes CDH2 and Vimentin as assessed via qRT-PCR and reported in **Fig. 3 G**.

### Reprogrammed mammary cancer cells overexpress TSPO

To further study the role of TSPO in the maladaptive conduit which exploits cell signalling to favour malignant transformation we devised isogenic lines with acquired endocrine therapy resistance via different mechanisms. CF41 cells were cultured with 4-OH TAM (5 μM) for 12 months to gain an *in vitro* model of tamoxifen-resistant cells (CF41-TAM) (i) or kept for an equal length of time in a hormone-deprived medium (CF41-LTHD) to mimic a model of endocrine manipulation and aromatase inhibitor resistance (ii) (**Fig. 4 A**). Reprogramming via Tamoxifen confirmed the upregulation of TSPO in CF41-TAM cells (**Fig. 4 B**). CF41-TAM cells not only overexpress TSPO but show altered expression of epithelial-to-mesenchymal transition (EMT) hallmark genes (**Fig. 4 C**). The susceptibility of both the newly established lines to cell death was consequently assessed in a variety of cytotoxic treatments to corroborate the protection enacted by TSPO. Both CF41-LTHD and CF41-TAM show reduced sensitivity to STS even though this was significantly lower in CF41-TAM cells (**Fig. 4 D**) whose level of mitochondrial cholesterol doesn’t change in response to 4-OH TAM (**Fig. 4 E,F**). CF41, CF41-TAM and CF41-LTHD were all exposed to 4-OH TAM and the TSPO ligand endoxifen[33] (**Fig. 4 G**). The latter showed greater potency and efficacy than 4-OH TAM in CF-41 cells control (i) (**Fig. 4 H**); greater potency in CF41-LTHD cells (ii) (**Fig. 4 I**); but none of the two in the CF41-TAM cells (iii) (**Fig. 4 J**).

**Figure 4.**
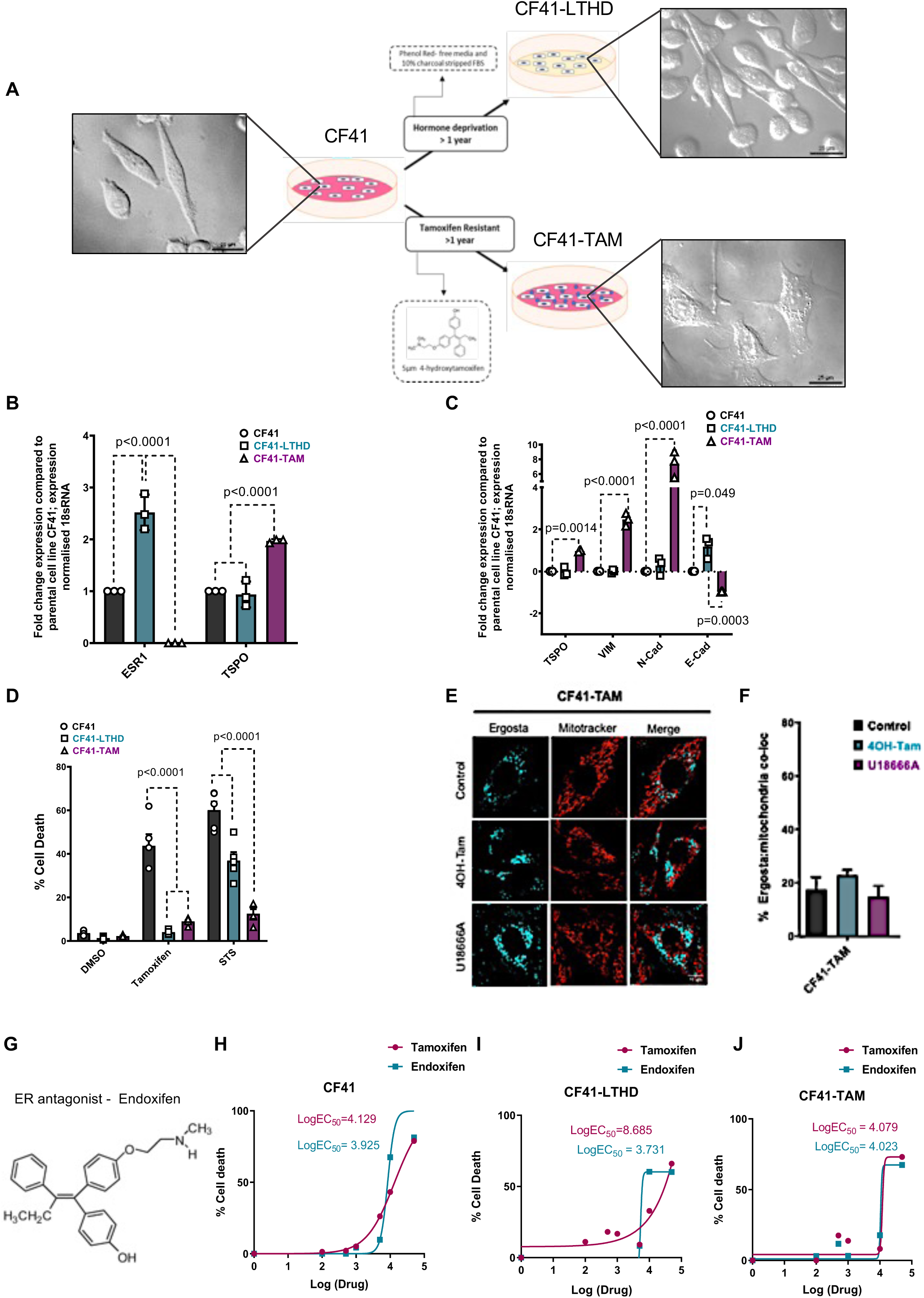
CMTs gain resistance to Tamoxifen and Oestrogen deprivation by upregulating TSPO. **A)** Schematic depicting the development of oestrogen-independent in vitro canine models of mammary cancer. CF41 cells were either deprived of hormones or treated with 5μM 4-hydroxytamoxifen for >1 year. Brightfield images of the cell lines were taken using differential interference contrast microscopy to show morphological changes. Scale bar = 25 μm. **B, C**) Gene expression analyses reveal a correlative relationship between upregulated mitochondrial TSPO and the expression of EMT genes in CF41-TAM cells. Fold change expression compared to parental cell line CF41, gene expression normalised to housekeeping gene 18sRNA (TSPO: CF41=0, CF41-LTHD 0.09±-0.151, CF41-TAM 0.99±0.95; VIM (vimentin): CF41=0, CF41-LTHD 0.05±0.05, CF41-TAM 2.64±2.30; CDH2 (N-cadherin): CF41=0, CF41-LTHD 0.51±0.03, CF41-TAM 8.45±6.39; CDH1 (E-cadherin): CF41=0, CF41-LTHD 1.47±0.88, CF41-TAM -0.97±-0.99; n=3). **D)** Percent cell death of oestrogen independent cell lines compared to parental CF41 following treatment with 10 μM 4-hydroxytamoxifen and staurosporine (STS) for 16h. DMSO treatment was used as a negative control. (DMSO: CF41 4.08±3.20, CF41-LTHD 1.61±1.00, CF41-TAM 2.48±2.02; Tamoxifen: CF41 49.0±38.4; CF41-LTHD 4.49±3.62, CF41-TAM 9.71±8.30; STS: CF41 64.0±56.2, CF41-LTHD 40.8±33.0; CF41-TAM 14.5±10.5; n=4). **E)** Confocal images of CF41-TAM cells loaded with mitotracker (red) and Ergosta (cyan) to stain mitochondria and cholesterol respectively following treatments with 4OH-tamoxifen and cholesterol trafficking inhibitor U18666A (0.1 µM9. **F**) Quantification of Ergosta (cholesterol analogue) co-localising with mitotracker following tamoxifen treatment in CF41-TAM cells (CF41-TAM Ctrl 22.29±13.08, 4OH-TAM 24.92±21.07; U18666A 18.97±10.90, 15 images were analysed per condition). **G)** Chemical structures of the active metabolite N-desmethyl-4-hydroxytamoxifen (Endoxifen). **H**) EC50 Log curve representing the dose (or concentration) causing 50% of cell death for Tamoxifen and Endoxifen in CF41 cells; n=3. **I)** EC50 Log curve representing the dose (or concentration) causing 50% of cell death for Tamoxifen and Endoxifen in CF41-LTHD cells; n=3. **J**) EC50 Log curve representing the dose (or concentration) causing 50% of cell death for Tamoxifen and Endoxifen in CF41-Tamoxifen cells; n=3.

Based on this we therefore propose a model in **Fig. 5 C** in which we highlight the level of mitochondrial cholesterol as discriminant in aggressive reprogramming as well as a target to counteract cellular proliferation.

**Figure 5.**
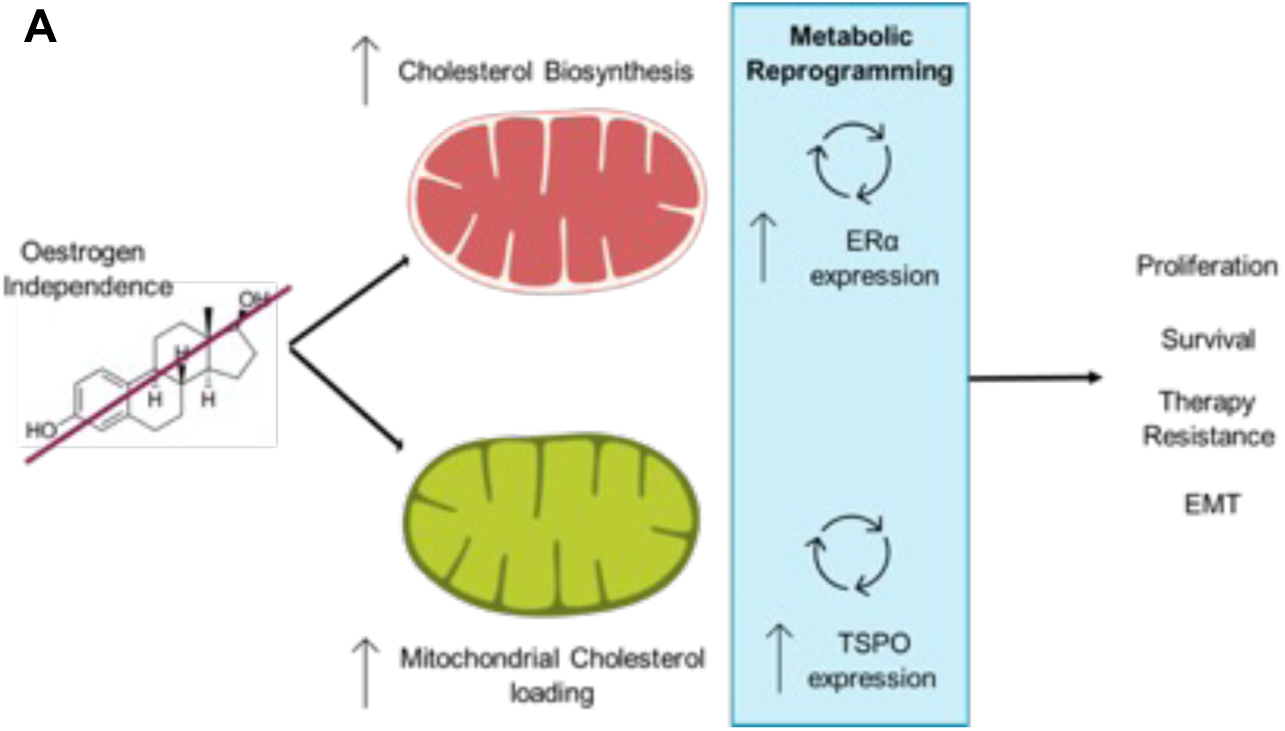
Proposed model for the TSPO mediated rewiring of mitochondrial cholesterol metabolism in CMTs. **A)** Working Model of the envisaged role played by mitochondrial cholesterol in cancer cells reprogramming to aggressive phenotype resistant to treatment.

## Discussion

Alterations in cholesterol uptake, metabolism, and homeostasis partake in the pathogenesis of human breast cancer[34]. Here we show that such a feature is retained in CMTs which exploit the 18kDa mitochondrial protein TSPO. TSPO, which is involved in the machinery of the cholesterol import from the cytosolic environment, is positively associated with the aggressiveness of canine mammary tumour tissues (**Fig. 1 A-C**) and cell lines[21] (**Fig. 2 A-D**).

Interestingly, the mRNA messenger coding for TSPO increases in samples of neutered dogs (**Fig. 1 F**). This common practice aimed at reducing the risk of tumour formation in female dogs^6^ does seem to engage TSPO hence the mitochondrial pathway for the handling of cholesterol. However, the TSPO pattern diverges from one of the Estrogen Receptors as highlighted by previous work which has reported that ESR1 (encoding for ERα) does not increase in malignant CMTs[11]. Similarly, the expression of ESR2 (encoding for ERβ) was found to be undetectable in most of the isolated samples from both healthy and diseased subjects[11]. The immunohistochemical data here shown (**Fig. 1 B, D**) are therefore in line with previous literature[11] by confirming that the aggressive lesions do not increase their ERβ over benign tumours. Moreover, the TSPO trend of upregulation according to malignancy matches what happens in the mammary type of tumour[29].

We are therefore tempted to speculate that the collapse of the natural circuit of hormone homeostasis, following neutering, triggers an intracellular response of adaptation which is read by TSPO upregulation. A response which predisposes to the production of steroids for which mitochondrial cholesterol is indispensable. However, the concomitant low level of EMT drivers (e.g. CDH-1) recorded in the same samples (**Fig. 1 G**) indicates that further events are required to trigger the neoplastic transformation and aggressive degeneration.

In human-derived breast cancer cells, we have previously demonstrated that TSPO-dependent modulation of cholesterol is essential for the mitochondrial retrograde response (MRR) which facilitates the nuclear localisation of the transcription factor NF-kB[29].

MRR is facilitated by points of contact, termed Nucleus Associated Mitochondria (NAM) which bridge mitochondria and the nucleus. The NAM molecular complex requires TSPO[31] on the outer mitochondrial membrane (OMM) to form. Repression of TSPO is consequently a suitable way to impair mitochondrial coupling with the nucleus and NF-kB function[29]. Here, we show that in canine cancer cells, the patterns of NF-kB are equally dependent on TSPO whose repression impairs the physiology of this transcription factor.

Modified NF-kB alters therefore the expression of pro-survival and oncogenic genes which dictate the proliferative capacity and resistance to anti-cancer treatment.

It was therefore not surprising that the pharmacological manipulation of TSPO significantly prevented CMT cells’ spherogenesis (i) (**Fig. 2 C-H**), migratory capacity (ii) (**SFig. 2 A, B**), susceptibility to chemically induced cell death (iii) (**Fig. 4 G-J**), and expression of the EMT genes associated with the development of cancer resistance to chemotherapy[35 36] (iv) (**Fig. 3 F, G**).

It should be also noted that the interdependence between spherogenesis and MRR is further confirmed by the data attained with the P62-mediated mitophagy inducer (PMI)[32] which by promoting autophagic clearance of mitochondria (mitophagy) would curtail the communication with the nucleus (**Fig. 2 I, J**). This confirms that mitochondrial accumulation is a determinant for the activation and execution of the adaptive signalling exploited by cancer cells[12].

Therefore, TSPO by counteracting mitophagy favours cellular adaptation to pathogenic cues[12] as well as driving resistance to chemotherapy in canine cells as well as in the human counterpart. Neutering (i), oestrogen deprivation (ii) and long-term exposure to 4-hydroxytamoxifen (4-OH TAM) (iii) all lead to an overexpression of TSPO (**Fig. 1 F**, **Fig. 4 A-C**): the required modification on mitochondria to facilitate their interplay with the nucleus.

A deeper understating of TSPO molecular function may therefore explain the lack of ET efficacy in canine patients in which the already high level of TSPO further increases in response to the selective oestrogen modulator favouring the aggressive proliferation of cells[29].

As confirmation of this, 4-OH TAM promotes cholesterol loading into the mitochondria and the high endogenous level of TSPO recorded in REM-134 cells increases the degree of perturbation further if compared to the initial basal levels of cholesterol (**Fig. 3 C, D; SFig. 2 C, D**). This kinetic may be also contributed by potential mutations in TSPO and the bigger mitochondrial network associated with TSPO level overexpression[22].

The efficacy of cellular susceptibility to chemically induced demise is dependent on TSPO as both the reprogrammed CF41 (CF41-TAM) and REM-134 cells, which express greater levels of the protein, are more resistant to STS and 4-OH TAM-mediated demise (**Fig. 4 D**). This is nonetheless reverted by TSPO pharmacological regulators (**Fig. 4 G-J**) mimicking what was previously seen in human cells[37].

In keeping with this, the TSPO ligand PK11195 -acknowledged to bind and repress TSPO- is also able to counteract the cancer stem cell subpopulation (**Fig. 2 I, J**) and migratory capacity (**SFigure 2 A, B**). All promising indications advocate for the suitability of TSPO ligands as co-adjuvants in therapeutic protocols in CMT cells.

In conclusion, TSPO is a molecule involved in the mitochondrial mechanisms of cholesterol import which holds etiopathogenic relevance in the progression of canine mammary tumours.

Modulation of TSPO expression induced by endocrine chemotherapeutics associates with a redistribution of cholesterol which we propose as a priming signature in this disease.

In canine cells, TSPO is therefore able to define a conduit of adaptative signalling and mito-nuclear communication which triggers genomic and epigenomic responses[38].

The side effects which currently limit -if not discourage- the use of endocrine chemotherapy in canine patients[11] may therefore have in TSPO a factor as well as a means for their understanding.

## Patent

No patents or patent applications have been derived from this work.

## Author Contributions

Conceptualization: L.H. and M.C.; Supervision: L.H. and M.C.; Methodology: L.H. G. B-H., B.K. M.R. and M.C.; Tissue collection, processing and tumour staging, knowledge transfer: M.P.K, I.M.R, F.G.; Investigation & Data Collection: L.H. G. B-H., B.K. M.R. and M.C.; Writing - Original Draft: L.H. and M.C.; Writing – Review & Editing: L.H., R.L.P., M.P.K, B.K., M.R. and M.C.

## Institutional Review Board Statement

The study was conducted by the standards of correct research practice adopted at the hosting institutions. All owners consented to their animal becoming a research participant and animal experimentation was approved by the Cantonal Veterinary Authority in Zurich, Switzerland (permission No. 136/2009 and 165/2012).

## Informed Consent Statement

Not Applicable

## Data Availability Statement

The data that support the findings of this study are available from the corresponding author upon reasonable request and they are stored in MCP laboratory data repositories at https://intranet.rvc.ac.uk/gdrive.

## Acknowledgements

We are grateful to Dr. Hibbert, Mr Simboy and the other dedicated lab technicians at the Royal Veterinary College who have made this work possible. CF35 cells were provided by Prof. Marilène Paquet (University of Montreal, Canada).

## References

1. Harbeck N, Penault-Llorca F, Cortes J, et al. Breast cancer. Nature Reviews Disease Primers 2019;5(1) doi: 10.1038/s41572-019-0111-2.

2. McCowan C, Shearer J, Donnan PT, et al. Cohort study examining tamoxifen adherence and its relationship to mortality in women with breast cancer. British Journal of Cancer 2008;99:1763–68 doi: 10.1038/sj.bjc.6604758.

3. Hultsch S, Kankainen M, Paavolainen L, et al. Association of tamoxifen resistance and lipid reprogramming in breast cancer. BMC Cancer 2018;18(1) doi: 10.1186/s12885-018-4757-z.

4. Ring A, Dowsett M. Mechanisms of tamoxifen resistance. Endocrine-Related Cancer, 2004:643–58.

5. Schneider R, Dorn CR, Taylor DON. Factors Influencing Canine Mammary Can-cer Development and Postsurgical Survival 1.2.

6. Beauvais W, Cardwell JM, Brodbelt DC. The effect of neutering on the risk of mammary tumours in dogs - a systematic review. Journal of Small Animal Practice 2012;53(6):314–22 doi: 10.1111/J.1748-5827.2011.01220.X.

7. Abdelmegeed SM, Mohammed S. Canine mammary tumors as a model for human disease (Review). Oncology Letters: Spandidos Publications, 2018:8195–205.

8. Holen I, Speirs V, Morrissey B, Blyth K. In vivo models in breast cancer research: progress, challenges and future directions. 2017 doi: 10.1242/dmm.028274.

9. Ghoncheh M, Pournamdar Z, Salehiniya H. Incidence and mortality and epidemiology of breast cancer in the world. Asian Pacific Journal of Cancer Prevention 2016;17:43–46 doi: 10.7314/APJCP.2016.17.S3.43.

10. Gazdar AF, Kurvari V, Virmani A, et al. CHARACTERIZATION OF PAIRED TUMOR AND NON-TUMOR CELL LINES ESTABLISHED FROM PATIENTS WITH BREAST CANCER. Int. J. Cancer 1998;78:766–74 doi: 10.1002/(SICI)1097-0215(19981209)78:6.

11. Spoerri M, Guscetti F, Hartnack S, et al. Endocrine control of canine mammary neoplasms: Serum reproductive hormone levels and tissue expression of steroid hormone, prolactin and growth hormone receptors. BMC Veterinary Research 2015;11(1) doi: 10.1186/s12917-015-0546-y.

12. Strobbe D, Sharma S, Campanella M. Links between mitochondrial retrograde response and mitophagy in pathogenic cell signalling. Cellular and Molecular Life Sciences 2021;78:3767–75 doi: 10.1007/s00018-021-03770-5.

13. Jeng M-H, Shupnik MA, Bender TP, et al. Estrogen Receptor Expression and Function in Long-Term Estrogen-Deprived Human Breast Cancer Cells*. 1998.

14. Santen RJ, Lobenhofer EK, Afshari CA, Bao Y, Song RX. Adaptation of estrogen-regulated genes in long-term estradiol deprived MCF-7 breast cancer cells. doi: 10.1007/s10549-005-5776-4.

15. Canadas-Sousa A, Santos M, Leal B, Medeiros R, Dias-Pereira P. Estrogen receptors genotypes and canine mammary neoplasia. BMC Veterinary Research 2019;15(1) doi: 10.1186/s12917-019-2062-y.

16. Estrogen metabolism and breast cancer: A risk model. Annals of the New York Academy of Sciences; 2009. Blackwell Publishing Inc.

17. Santen R, Cavalieri E, Rogan E, et al. Estrogen Mediation of Breast Tumor Formation Involves Estrogen Receptor-Dependent, as Well as Independent, Genotoxic Effects. doi: 10.1111/j.1749-6632.2008.03685.x.

18. Chen F-P, Chien M-H, Chen H-Y, Huang T-S, Ng Y-T. Effects of estradiol and progestogens on human breast cells: Regulation of sex steroid receptors. doi: 10.1016/j.tjog.2012.09.038.

19. Morris JS, Dobson JM, Bostock DE. Use of tamoxifen in the control of canine mammary neoplasia. Veterinary Record 1993;133(22):539–42 doi: 10.1136/VR.133.22.539.

20. Tavares WLF, Lavalle GE, Figueiredo MS, et al. Evaluation of adverse effects in tamoxifen exposed healthy female dogs. Acta Veterinaria Scandinavica 2010;52(1) doi: 10.1186/1751-0147-52-67.

21. Denora N, Natile G. Molecular Sciences Editorial An Updated View of Translocator Protein (TSPO). doi: 10.3390/ijms18122640.

22. Gatliff J, Campanella M. TSPO: kaleidoscopic 18-kDa amid biochemical pharmacology, control and targeting of mitochondria. Biochemical Journal 2016;473(2):107–21 doi: 10.1042/BJ20150899.

23. Galano M, Li Y, Li L, Sottas C, Papadopoulos V. Role of Constitutive STAR in Leydig Cells. International Journal of Molecular Sciences 2021, Vol. 22, Page 2021 2021;22(4):2021–21 doi: 10.3390/IJMS22042021.

24. Rone MB, Midzak AS, Issop L, et al. Identification of a Dynamic Mitochondrial Protein Complex Driving Cholesterol Import, Trafficking, and Metabolism to Steroid Hormones. Molecular Endocrinology 2012;26:1868–82 doi: 10.1210/me.2012-1159.

25. Han Z, Slack RS, Li W, Papadopoulos V. Expression of Peripheral Benzodiazepine Receptor (PBR) in Human Tumors: Relationship to Breast, Colorectal, and Prostate Tumor Progression. Journal of Receptors and Signal Transduction 2003;23(2-3):225–38 doi: 10.1081/RRS-120025210.

26. Rechichi M, Salvetti A, Chelli B, et al. TSPO over-expression increases motility, transmigration and proliferation properties of C6 rat glioma cells. 2007 doi: 10.1016/j.bbadis.2007.12.001.

27. Bhoola NH, Mbita Z, Hull R, Dlamini Z. Translocator Protein (TSPO) as a Potential Biomarker in Human Cancers. International Journal of Molecular Sciences Article doi: 10.3390/ijms19082176.

28. Shaiken TE, Opekun AR. Dissecting the cell to nucleus, perinucleus and cytosol There are amendments to this paper. 2014 doi: 10.1038/srep04923.

29. Desai R, East DA, Hardy L, et al. Mitochondria form contact sites with the nucleus to couple prosurvival retrograde response. Sci. Adv, 2020:9955–73.

30. Feoktistova M, Geserick P, Leverkus M. Crystal Violet Assay for Determining Viability of Cultured Cells. Cold Spring Harbor Protocols 2016;2016(4):pdb.prot087379-pdb.prot79 doi: 10.1101/PDB.PROT087379.

31. Shaw FL, Harrison H, Spence K, et al. A Detailed Mammosphere Assay Protocol for the Quantification of Breast Stem Cell Activity. doi: 10.1007/s10911-012-9255-3.

32. McIntosh AL, Atshaves BP, Huang H, Gallegos AM, Kier AB, Schroeder F. Fluorescence techniques using dehydroergosterol to study cholesterol trafficking. Lipids, 2008:1185–208.

33. Ma X-J, Wang Z, Ryan PD, et al. A two-gene expression ratio predicts clinical outcome in breast cancer patients treated with tamoxifen.

34. East DA, Fagiani F, Crosby J, et al. Chemistry & Biology Resource PMI: A DJ m Independent Pharmacological Regulator of Mitophagy. doi: 10.1016/j.chembiol.2014.09.019.

35. Strobbe D, Campanella M. Anxiolytic Therapy: A Paradigm of Successful Mitochondrial Pharmacology. doi: 10.1016/j.tips.2018.02.008.

36. Kuzu OF, Noory MA, Robertson GP. The Role of Cholesterol in Cancer. 2016 doi: 10.1158/0008-5472.CAN-15-2613.

37. Campanella M. The Physiology and Pharmacology of the Mitochondrial 18 kDa Translocator Protein (TSPO): An Emerging Molecular Target for Diagnosis and Therapy. Editorial Current Molecular Medicine, 2012:355–55.

38. Galle E, Thienpont B, Cappuyns S, et al. DNA methylation-driven EMT is a common mechanism of resistance to various therapeutic agents in cancer. doi: 10.1186/s13148-020-0821-z.

