## Supplementary Figures for "Canine Mammary Tumours (CMTs) exploit Mitochondrial Cholesterol for aggressive reprogramming"

#### Supplementary Figure 1

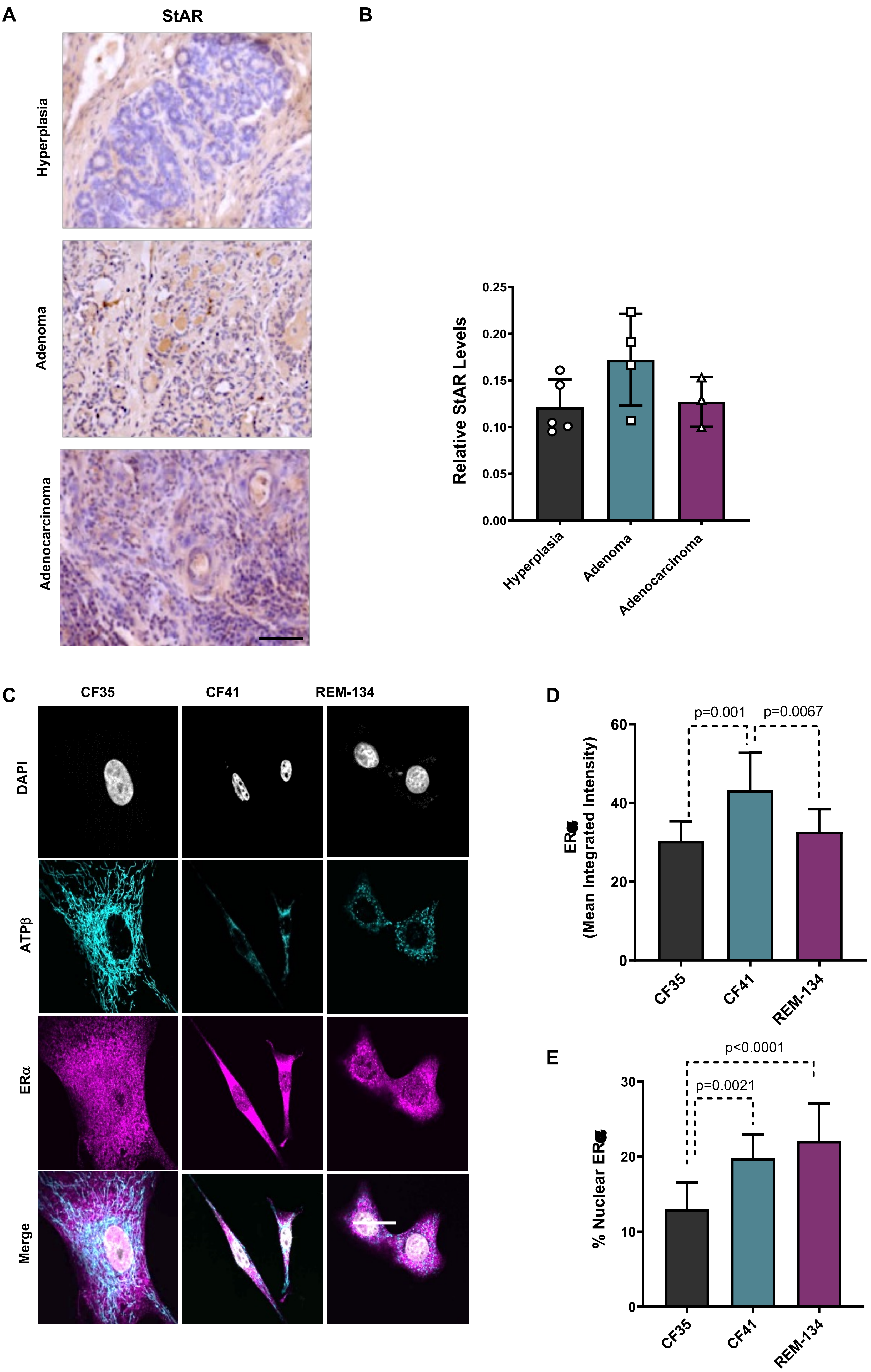

### Supplementary Figure 2

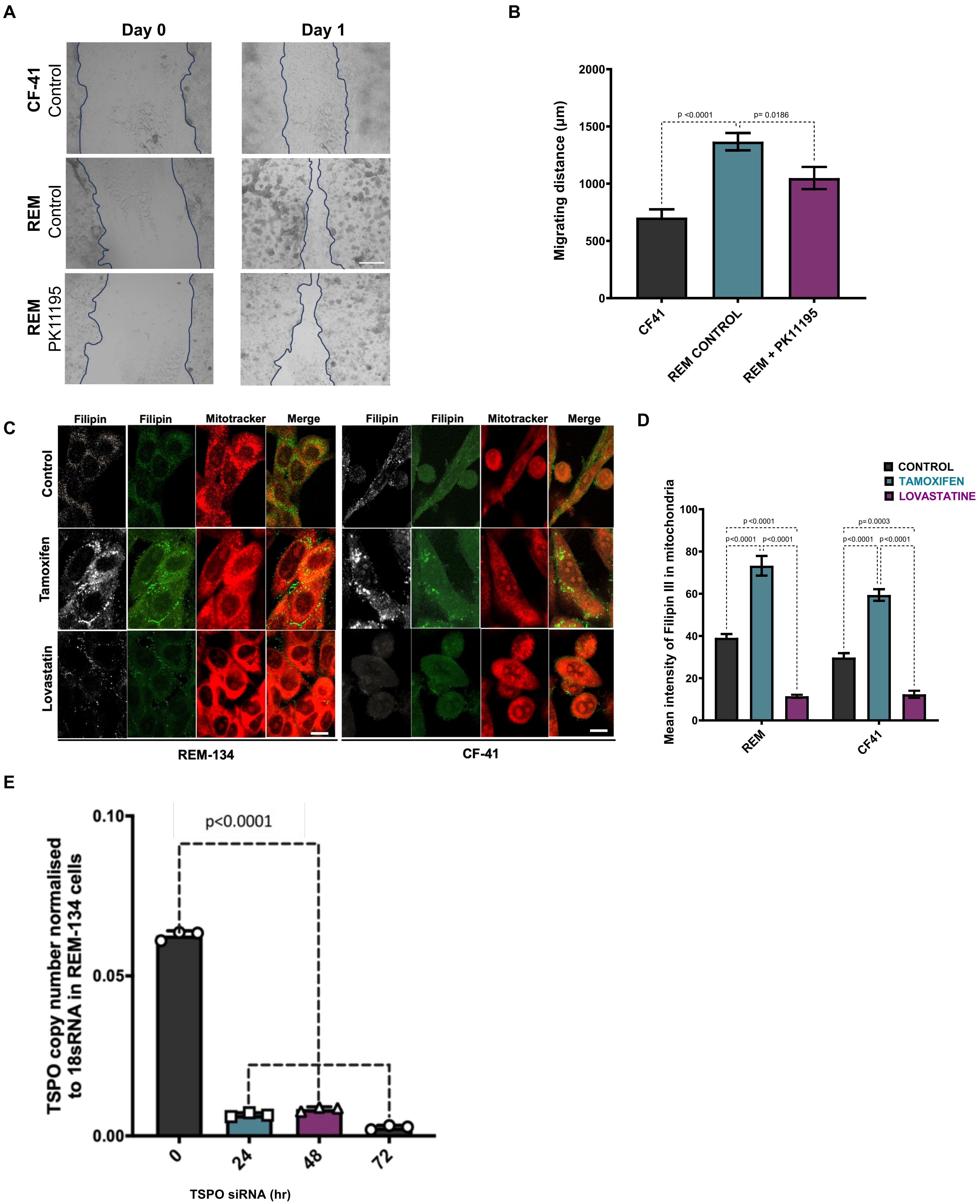

**Supplementary Figure 1. STAR expression profile in canine breast cancer tissues and Pattern of ERa in canine mammary tumour cells.**

**A)** Immunohistochemical staining of primary canine breast cancer tissue with varying pathophysiological grades labelled with StAR antibody visualized using DAB (brown) and co-stained with haematoxylin (purple). Scale bar=100um. **B)** Quantification of StAR levels in primary canine mammary tumours at different pathohistological grades. (Hyperplasia 0.15± 0.09; Adenoma 0.22 ± 0.12; Adenocarcinoma 0.15 ± 0.10; n=5, p>0.05 ns). **C**) Immunocytochemical analysis of ERa receptors in CF35, CF41 and REM-134 cells quantified in **D** and **E**. Scale bar= 10um.

**Supplementary Figure 2. Proliferative potential, cholesterol dynamics and chemical-induced demise in canine mammary tumour cells amenable to TSPO modulation.**

**A)** Representative images of the scratch-wound assay in CF-41, REM-134 cells and REM-134 cells co-treated with PK11195 (200nM); Scalebar = 500 µm and in **B** is reported the quantification of the distance covered by the cells in 24h, n=4.
**C**) Representative images of the Filipin and mitoTracker co-staining on both CF-41 and REM-134 cells challenged with Tamoxifen (10mM) and Lovastatin (10mM); Scalebar=25 µm. **D**) Quantification of the Filipin redistribution in the mitochondria (n=8). **E)** qPCR showing effective knockdown of TSPO using canine siRNA in REM-134 cells at 0, 24, 48 and 72h. TSPO expression normalized to 18sRNA (n=3).
