## Supplementary figures and images for "Canine Mammary Tumours (CMTs) exploit Mitochondrial Cholesterol for aggressive reprogramming"

### Tables

**Table 1**

RT-qPCR primers

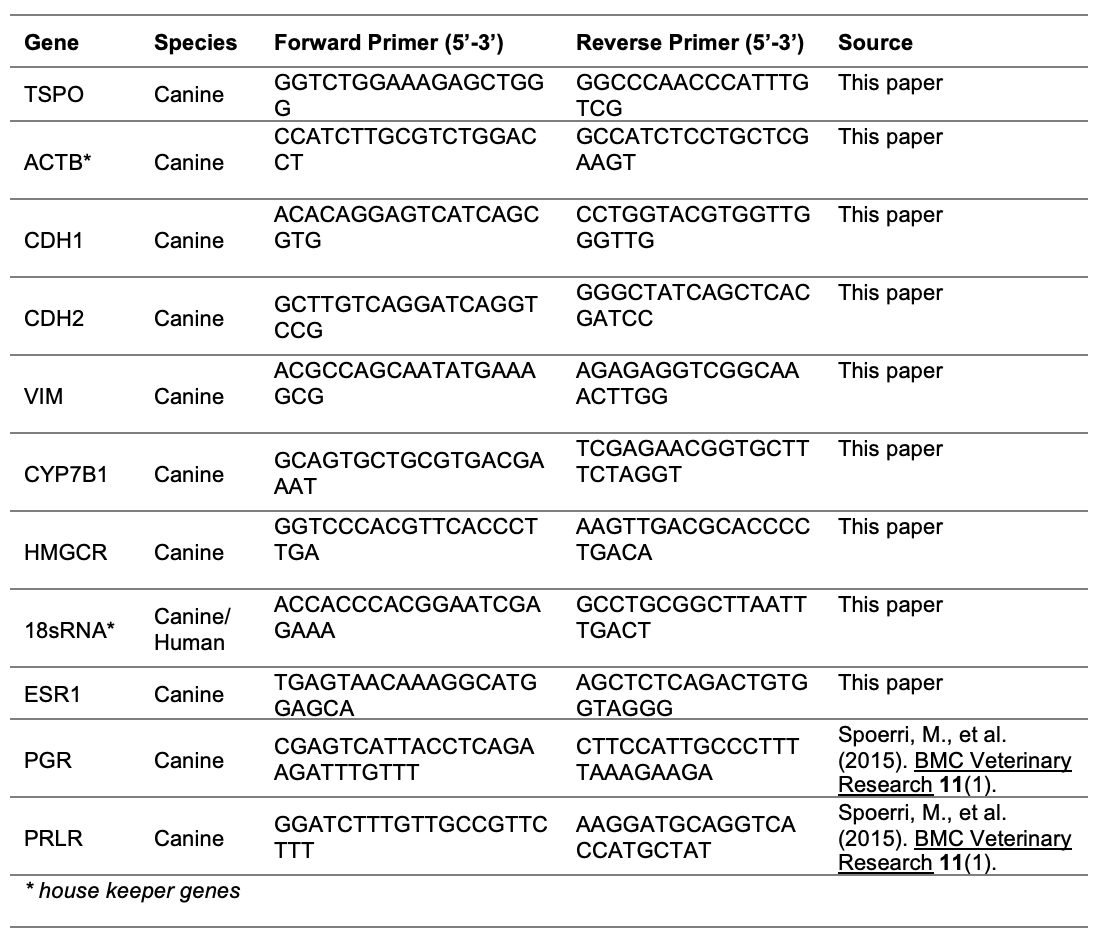

**Table 2**

qRT-PCR thermocycling conditions

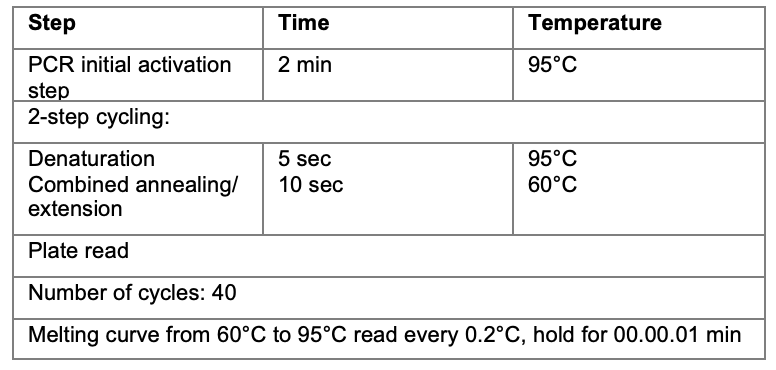
